# Training Infrastructure as a Service

**DOI:** 10.1101/2020.08.23.263509

**Authors:** Helena Rasche, Björn Grüning

**Author notes:** Contributed equally.

## Abstract

**Background:** Hands-on training, whether it is in Bioinformatics or other scientific domains, requires significant resources and knowledge to setup and run. Trainers must have access to infrastructure that can support the sudden spike in usage, with classes of 30 or more trainees simultaneously running resource intensive tools. For efficient classes, the jobs must run quickly, without queuing delays, lest they disrupt the timetable set out for the class. Often times this is achieved via running on a private server where there is no contention for the queue, and therefore no or minimal waiting time. However, this requires the teacher or trainer to have the technical knowledge to manage compute infrastructure, in addition to their didactic responsibilities. This presents significant burdens to potential training events, in terms of infrastructure cost, person-hours of preparation, technical knowledge, and available staff to manage such events.

**Findings:** Galaxy Europe has developed Training Infrastructure as a Service (TIaaS) which we provide to the scientific commnuity as a service built on top of the Galaxy Platform. Training event organisers request a training and Galaxy administrators can allocate private queues specifically for the training. Trainees are transparently placed in a private queue where their jobs run without delay. Trainers access the dashboard of the TIaaS Service and can remotely follow the progress of their trainees without in-person interactions.

**Conclusions:** TIaaS on Galaxy Europe provides reusable and fast infrastructure for Galaxy training. The instructor dashboard provides visibility into class progress, making in-person trainings more efficient and remote training possible. In the past 24 months, > 110 trainings with over 3000 trainees have used this infrastructure for training, across scientific domains, all enjoying the accessibility and reproducibility of Galaxy for training the next generation of bioinformaticians. TIaaS itself is an extension to Galaxy which can be deployed by any Galaxy administrator to provide similar benefits for their users. https://galaxyproject.eu/tiaas

## Findings

### Background

With the volume of bioinformatics data becoming available, the availability of training for bioinformaticians is not keeping up ([1]). The Galaxy platform [2] provides one such infrastructure on which to conduct trainings, it provides a user-friendly web-based interface to command line tools. With a wide range of tools across bioinformatics domains and beyond, and preexisting popularity within the life sciences community, it is an ideal platform for training ([3]).

The Galaxy community has developed a wide array of hands-on training materials covering the bioinformatics domains and beyond ([4]) in an attempt to combat this issue, but in order to run these tutorials at scale, one occasionally needs access to significant resources. The quite popular “Reference-based RNA-Seq data analysis” tutorial uses the STAR aligner ([5]), and while an ultra-fast aligner is ideal during training, it also consumes 32-64 GB of RAM at minimum. Running a single STAR job might be fine, but if 30 trainees run STAR simultaneously, it can present unexpected delays due to jobs contending for available compute resources. Most Galaxy servers submit jobs to compute clusters, but these clusters must also meet the needs of all of theother users of the Galaxy servers, and might not have enough resources for the requested jobs. Jobs thus queue and wait until resources become available.

Many tools across Galaxy have similarly large resource requirements, requirements which are only discoverable through testing. If trainers choose to manage their own infrastructure they need to obtain this knowledge via testing, and then configure their infrastructure to support the minimum memory and CPU requirements. Alternatively if the instructor chooses to use one of the many public Galaxy servers, they can rely on those services, but their trainees’ jobs could unexpectedly sit in shared queues for long periods of time. This is a significant barriers for potential, motivated trainers who wish to give a Galaxy or bioinformatics training courses.

We have developed Training Infrastructure as a Service (TIaaS) initially to make our own courses run more smoothly due to previous experiences with the aforementioned problems. TIaaS removes the main bottlenecks of training: setting up the training infrastructure is nearly completely automated, jobs are guaranteed to have private resources available so they can always run without delay, and it ensures separation of responsibilities between trainers who are teaching and the server administrators responsible for Galaxy. We have shared this service with the Galaxy training community to very positive feedback.

### Results

We implemented a Django based web service and job scheduling rules which function together to present a private queue for users in specific Galaxy user groups. This system can be generically useful for temporary and private compute resources, but is currently focused on training use case. Trainers register their course and teaching materials with an online form. Galaxy administrators review requests, using information about their class size, the tools used in those training materials, and the resource allocations of those tools on the infrastructure to estimate the required compute resources.

If space is available and any other criteria is met, the training can then be approved. Next, administrators optionally deploy private compute resources in a local cloud and attach them to their existing Galaxy scheduling. Administrators are provided with a URL such as https://usegalaxy.eu/join-training/test, which they share with the trainer, and the trainer can then in turn provide to their trainees. When training participants access the URL, they are registered in the TIaaS system, without the service administrator or trainers needing to be aware of users identities for optimal GDPR compliance. The scheduler, now aware of the training group, will grant any job run by someone in that group the ability to run on the private training nodes (Figure 2).

The course dashboard, visualising the progress of participants (Figure 1), has significantly eased the lives of trainers. The dashboard provides instantaneous, aggregated, and anonymized feedback for the trainers into how their trainees are progressing in a GDPR compliant manner. It has also enabled remote and hybrid trainings to take place, where trainers present a topic via video and it is broadcast to numerous individuals or sites. Requiring students or teaching sites to constantly provide up-to-date information on how participants are progressing simply does not work. With the training dashboard this is unnecessary as trainers, helpers, and even trainees can see how everyone is progressing online, instantaneously. Teachers can see if steps are completed successfully on average, or if there are many issues and they might need to pause or explain the step in more detail.

**Figure 1.**
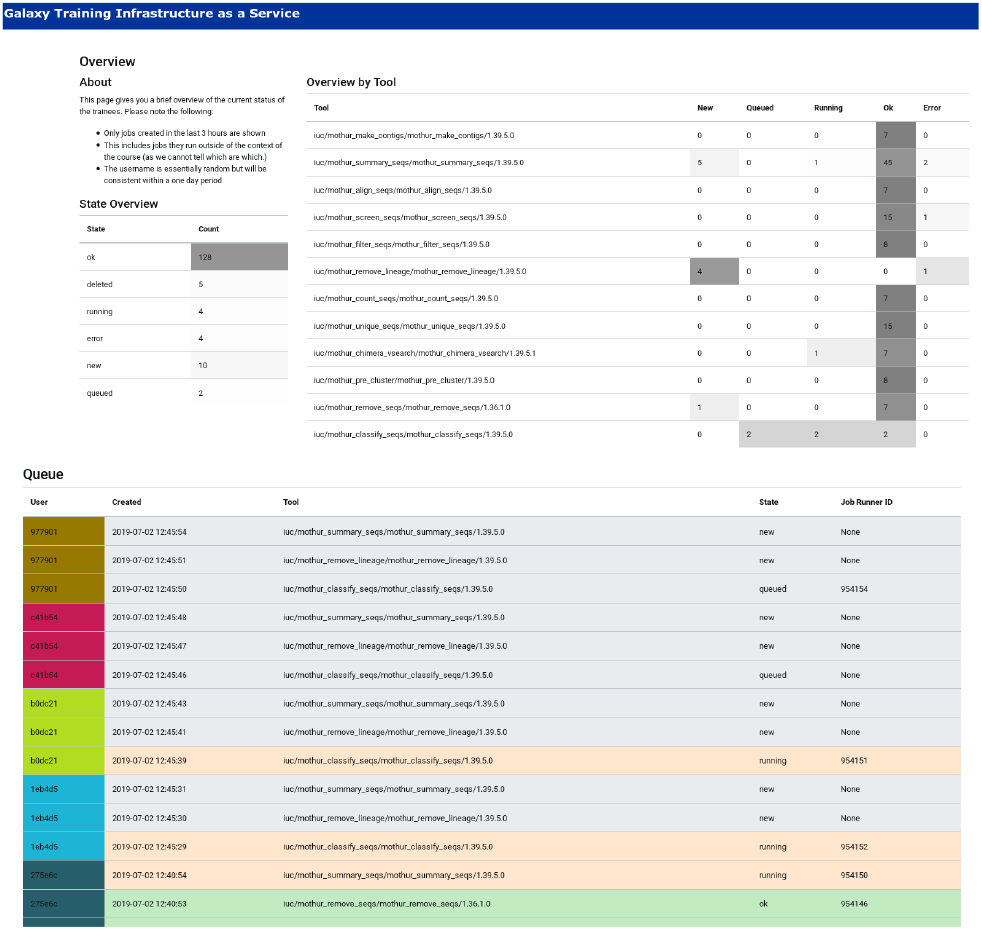
The top of the training dashboard page shows the status of the jobs in the past hours. A heatmap of the tools which were run indiciates if everything is running smoothly or if there is anything the trainer should look into. As trainees follow along and run different tools these show up immediately, allowing trainers to identify if everyone has started or finished a specific step.. The bottom image shows the rest of the training dashboard, which lists jobs that were run chronologically, colour coded first by user, and second by the job status. Random identifiers are used to protect user privacy as we wanted dashboards to be public.

**Figure 2.**
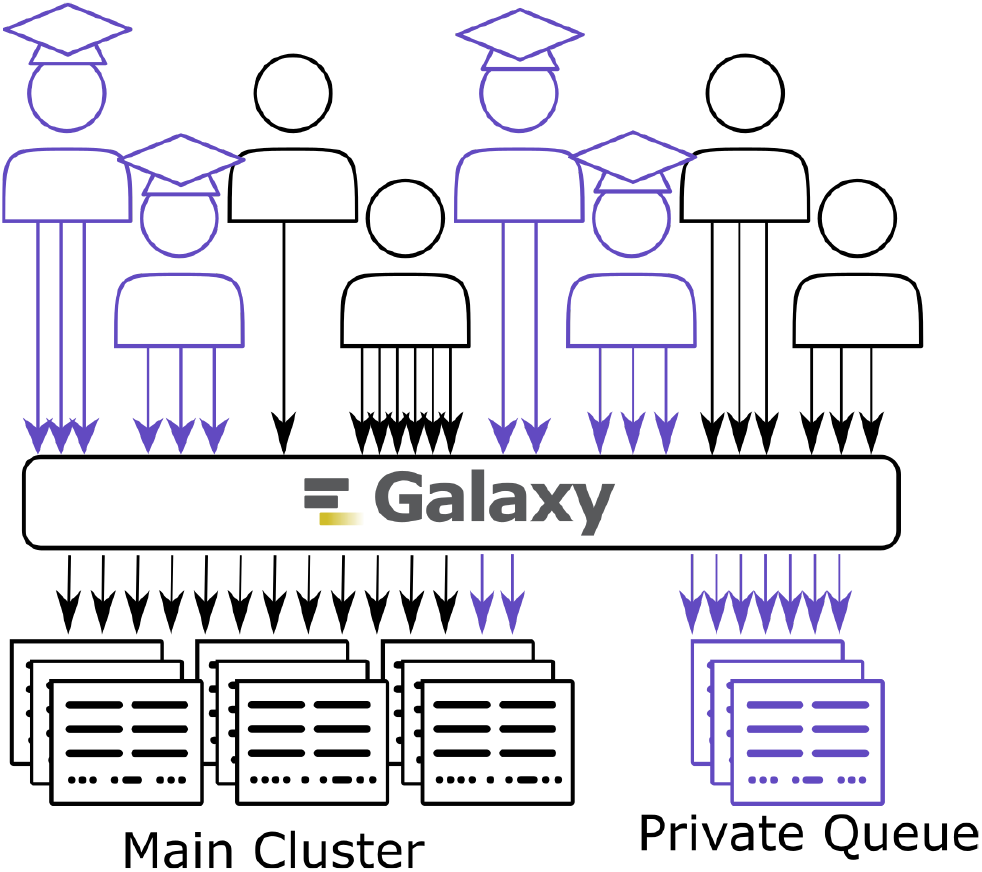
All jobs are processed by the same Galaxy server, but jobs from users in the training groups receive special handling. These jobs are allowed to run on the protected training resources (purple). If the training resource is full, these jobs can spill over to the main queue if necessary.

#### Usage

Since the introduction of TIaaS in July 2018, it has seen nearly constant use with more than 110 trainings occuring on the platform, all across the world (Figure 3). Everything from one day workshops for bioinformaticians to multi-month courses for highschool students have been hosted by TIaaS and UseGalaxy.eu, covering topics such as HTS, RNA-Seq, ChlP-Seq, Imaging Analysis, Proteomics, and Machine Learning.

**Figure 3.**
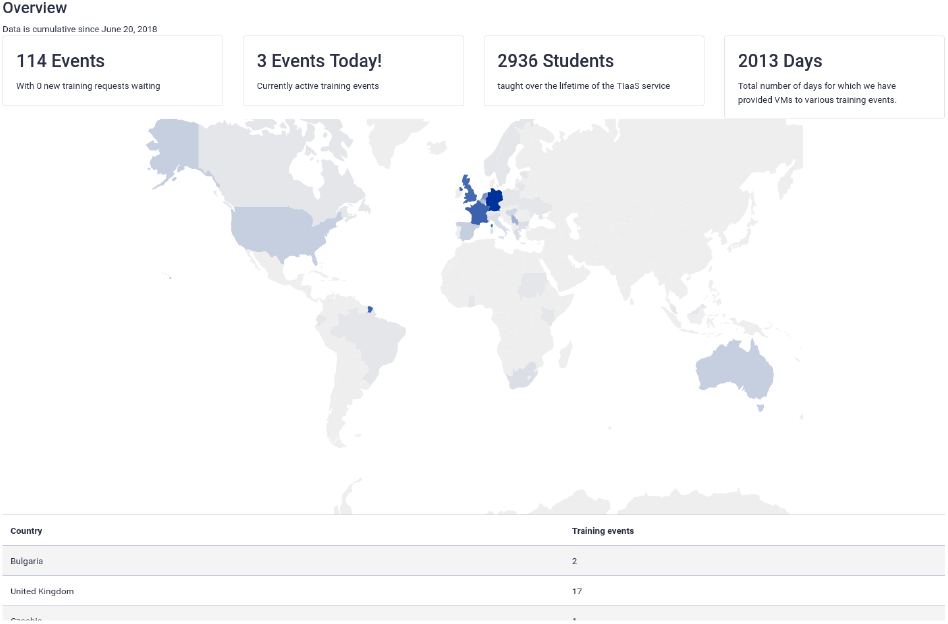
Since its introduction, TIaaS has seen worldwide usage despite-or perhaps in some cases due to-the server being hosted in Germany. This dashboard is part of the TIaaS platform, allowing deployers to share their numbers with funding agencies and the world.

Class sizes have ranged considerably, from the median of 15 participants to a maximum of 80 at a Transcriptomics workshop. Most course were quite short training events with a median of two days, however some ran for multiple months like a highschool course that used the service for three months, over the course of a semester, teaching Metagenomics. As the system is quite generic, it can easily accomodate a range of use cases.

## Methods

### Implementation

TIaaS was written in Python with the Django framework. It has been designed from the start to have a very limited scope: provide a form to register events, an approval flow for administrators, and a connection to the Galaxy database to manage user-group relationships when needed.

For instructors a form is provided (`/tiaas/new`) which permits them to register a new training event. When submitted, this goes into the associated database. Administrators can view the requested training events and approve or reject them using the built in Django admin interface. When users visit their their training url (`/join-training/<id>`) the system accesses their Galaxy session cookie, and decodes it. They are automatically registered as part of a Galaxy group named after the training (e.g. `training-<id>`) which is created as needed.

When instructors or students visit the dashboard (Figure 1), the training ID is extracted from the URL (e.g. “test” from https://usegalaxy.eu/join-training/test/status), and all nonterminal jobs, in the past 1-6 hours, from those users are presented in a de-identified manner.

When a job is submitted by a user in a training group, UseGalaxy.eu’s job scheduling system system reads the user’s groups and roles, and if any of these include something pre-fixed with `training-`, then this is converted to a job scheduler specific requirement string (Figure 4). Ideally these are constructured to prefer training nodes, and spill over to the main queue if training nodes are full, but this feature is dependant on specific scheduler capabilities.

**Figure 4.**
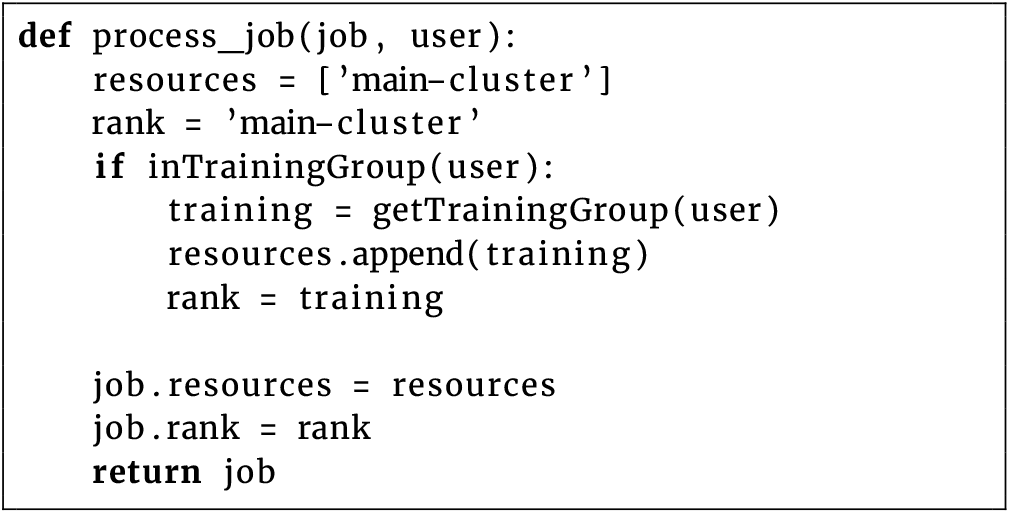
Python pseudocode representing how to process TIaaS jobs and allocate them to a private queue.

### Deploying

As the Galaxy community has largely settled on Ansible for deployment of Galaxy, and related components, an Ansible role was produced for deploying the TIaaS Service.

## Availability of source code and requirements

- Project name: Training Infrastructure as a Service
- Github repository: https://github.com/usegalaxy-eu/tiaas2/
- Training Manual: https://training.galaxyproject.org/training-material/topics/instructors/tutorials/setup-tiaas-for-training/tutorial.html
- Operating system(s): Unix
- Other requirements: Galaxy version 18.01 or higher
- License: GNU AGPL-3.0

## Availability of supporting data and materials

All code is open source and available on GitHub (https://github.com/usegalaxy-eu/tiaas2).

## Declarations

### List of abbreviations

TIaaS: Training Infrastructure as a Service

## Author’s Contributions

HR and BG conceived of the presented idea. HR carried out the implementation. HR and BG contributed to the writing of the manuscript.

## Acknowledgements

The authors would like to thank the Galaxy community for their enthusiasm for this project, and their feedback on each iteration.

## References

1. Attwood TK, Blackford S, Brazas MD, Davies A, Schneider MV. A global perspective on evolving bioinformatics and data science training needs. Briefings in Bioinformatics 2017 Aug;20(2):398–404. https://doi.org/10.1093/bib/bbx100.

2. Afgan E, Baker D, Batut B, Van Den Beek M, Bouvier D, Čech M, et al. The Galaxy platform for accessible, reproducible and collaborative biomedical analyses: 2018 update. Nucleic acids research 2018;46(W1):W537–W544.

3. Batut B, Hiltemann S, Bagnacani A, Baker D, Bhardwaj V, Blank C, et al. Community-Driven Data Analysis Training for Biology. Cell Systems 2018 Jun;6(6):752–758.e1. https://doi.org/10.1016/j.cels.2018.05.012.

4. Galaxy Training Materials;. https://training.galaxyproject.org.

5. Dobin A, Davis CA, Schlesinger F, Drenkow J, Zaleski C, Jha S, et al. STAR: ultrafast universal RNA-seq aligner. Bioinformatics 2012 Oct;29(1):15–21. https://doi.org/10.1093/bioinformatics/bts635.

